# Gene-culture co-inheritance of a behavioral trait

**DOI:** 10.1101/069286

**Authors:** Elliot G. Aguilar, Erol Akçay

## Abstract

Human behavioral traits are complex phenotypes that result from both genetic and cultural transmission. But different inheritance systems need not favor the same phenotypic outcome. What happens when there are conflicting selection forces in the two domains? To address this question, we derive a Price equation that incorporates both cultural and genetic inheritance of a phenotype where the effects of genes and culture are additive. We then use this equation to investigate whether a genetically maladaptive phenotype can evolve under dual transmission. We examine the special case of altruism using an illustrative model, and show that cultural selection can overcome genetic selection when the variance in culture is sufficiently high with respect to genes. Finally, we show how our basic result can be extended to non-additive effects models. We discuss the implications of our results for understanding the evolution of maladaptive behaviors.

## 1 Introduction

Behavioral traits are among the most complex phenotypes under study in evolutionary biology. At the heart of that complexity is the interaction between genetic inheritance and the environment (Turkheimer, 2000). In organisms with social learning, a significant component of the environment can be conspecific individuals who will serve as models for socially learned behaviors, leading to the emergence of cultural transmission. Thus, in organisms with significant social learning, some behaviors will be determined by both genetic and nongenetic inheritance systems. Examples of behaviors that are influenced by both genetic and cultural transmission span a wide range, such as antisocial behavior (Maes et al., 2007), parental behavior (Pérusse et al., 1994), handedness (Laland, 2008), fertility (Alvergne et al., 2011; Colleran and Mace, 2015; Colleran, 2016; Kosova et al., 2010), and Alzheimer’s disease biomarkers (Levy et al., 2016) in humans, as well as song form in passerine birds Feher et al. (2009); Freeberg (2000), mate choice in Trinidadian guppies (Dugatkin, 1992), and foraging behavior in bottlenose dolphins (Krŭtzen et al., 2005). In each of these cases, an important fitness-related trait or behavior is determined not only by genetic inheritance, but also cultural transmission from individuals that may not have contributed any genetic material. Future investigations are likely to reveal even more examples across behavioral domains and taxa. Given the likely importance of genetic and nongenetic inheritance in determining so many behavioral traits, it is imperative to develop a better theoretical understanding of how such co-inheritance affects the evolution of behavioral traits.

In recent years, evolutionary theorists have begun to investigate the consequences of multiple inheritance systems (Otto et al., 1995; Day and Bonduriansky, 2011; Bonduriansky and Day, 2009). In a pair of papers, Day & Bondurianski used the Price equation to construct a general framework for modeling genetic and nongenetic traits that jointly determined phenotype, though their method only kept track of the reproductive fitness consequences of both systems of inheritance. While this approach gives a mathematically valid description of evolutionary change in a trait, it obscures the separate roles of genetic and cultural inheritance and selection in causing that change. To account for these separate causal roles, one needs to consider fitness measures in both systems of inheritance. A fitness measure implies a mapping from ancestral to descendant individuals; ancestors who map to more descendants have higher fitness. Multiple inheritance systems mean the possibility of multiple mappings, as illustrated with the following example. Imagine a population of asexual organisms (as in Figure 1) with a phenotype *p* determined by genetic and cultural inheritance. Let *p_a_* be the phenotype of an ancestor and *p_d_* the phenotype of her genetic descendant. If both genes and culture are inherited from the same ancestor, and we assume no flaws in transmission, then *p_a_ = p_d_.* However, if one’s genetic parent and cultural role model are not the same individual, then it is possible that *p_a_ ≠ p_d_.* If we consider the mapping solely from genetic parents to offspring, this discrepancy will appear simply to be an unexplained deviation between parents and offspring. However, we might also keep track of the mapping between cultural role models and pupils. Under this mapping, we might find that certain individuals map to more cultural descendants as a result of their phenotype due to selection in the cultural domain. Thus what appears under one mapping to be an unexplained deviation between parent and offspring is revealed under another mapping to be a force of selection in its own right.

**Figure 1:**
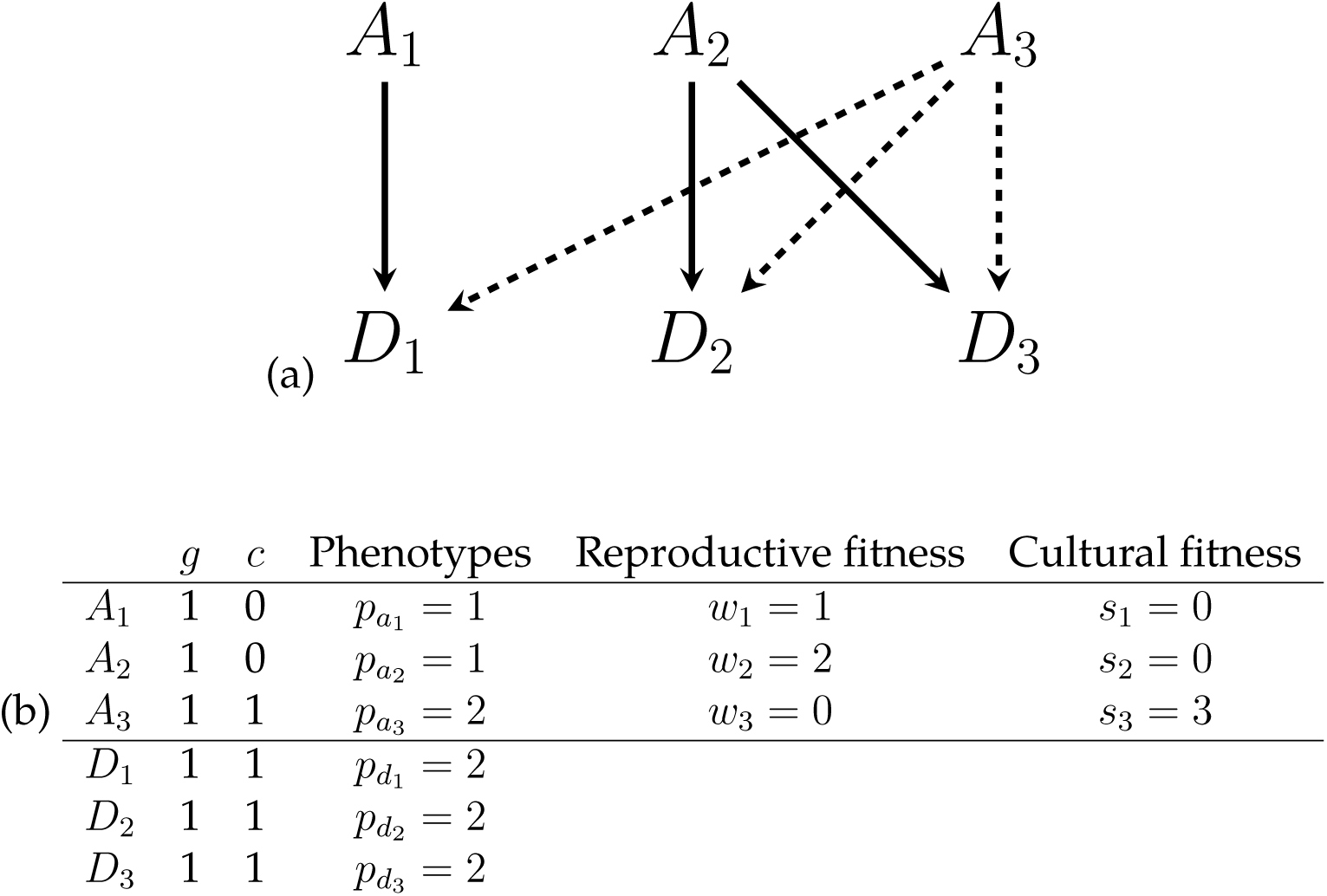
(a) The diagram shows the hereditary relationships between ancestors (*A*_1_,*A*_2_,*A*_3_) and descendants (*D*_1_,*D*_2_,*D*_3_). The solid line indicates reproductive relationships, while the *dashed* lines show cultural learning. While A_3_ sired no offspring, he is the cultural learning model for all descendants. (b) Gentotype, culture-type, phenotypic and fitness values for each ancestor and descendant (excepting fitness values). Each descendant has only one genetic and cultural ancestor, thus each solid edge corresponds to *v_ij_* = 1 and each dashed edge, *γ_ij_* = 1.

The argument above underscores the importance of considering fitness in each domain of inheritance when multiple forms of inheritance are present. We are not the first authors to highlight this point. Nearly forty years ago, Richerson & Boyd (1978) remarked that when both genes and culture determine a single phenotype, the value of the phenotype that maximizes genetic fitness may differ from the value that maximizes cultural fitness, leading to conflicts between the two inheritance systems. Using a weak selection model they showed that under certain conditions, the cultural fitness optimum could be reached at the expense of genetic fitness. In the ensuing decades, cultural evolution theory has largely focused on the case when the genetic trait encodes a learning rule that determines how a cultural trait is acquired (Boyd and Richerson, 1988; Cavalli-Sforza and Feldman, 1981; Boyd et al., 2003; Guzmán et al., 2007; Lehmann et al., 2008). By contrast, the problem of conflict between inheritance systems that affect the same trait has received surprisingly little attention, with the notable exception of the model of Findlay (Findlay, 1992), which only treated vertical cultural transmission. Indeed, some recent papers (El Mouden et al., 2014; Morin, 2014) claim that such conflicts will always be resolved in favor of reproductive fitness.

In this paper, we present a general approach to the evolution of a co-inherited trait by deriving a Price equation that explicitly incorporates both forms of inheritance. The Price equation is an exact description of an evolutionary process under a certain set of minimal assumptions (Price et al., 1970; Frank, 1998; Rice, 2004). Soon after its introduction, Hamilton (1975) pointed out that the Price equation can apply equally well to cultural transmission, and recent authors have developed it exclusively for that purpose (Henrich, 2004a; El Mouden et al., 2014). Others have also extended the Price equation to include multiple forms of inheritance (Day and Bonduriansky, 2011; Helanterä and Uller, 2010), though with the limitation of a single fitness measure. Here, we use a simple additive model to derive a Price equation that incorporates both domains of inheritance and their relevant fitness measures directly. We then analyze the condition for the evolution of a phenotype when selection in the two domains is in conflict. We take altruistic behaviors as a special case and present a series of illustrative models to explore the implications of our results, including assortative interactions and gene-culture correlations. Our model elucidates the conditions under which selection in one domain can overcome counter-selection in the other domain. We then extend our Price equation framework to more complicated models. We end with a discussion of the implications of our results for understanding the evolution of reproductively maladaptive behaviors.

## 2 Gene-Culture Price equation

We model the evolution of a trait that results from both genetic and cultural inheritance. Evolution here means the change in the phenotypes in a population, not only the change in the genetic or culturally inherited information that underlies them. An individual’s phenotype is represented by a continuous variable, *p*. We can take this to represent a behavioral trait, such as one of the big five personality traits (e.g. extraversion, agreeableness, conscientousness, etc.) (Goldberg, 1993). We assume that the effects of genetic and cultural inheritance are additive, i.e., we express an individual’s phenotype as the following:

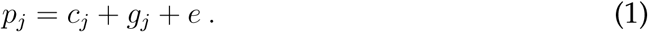

The final term, *e*, is the effect of the environment that does not include cultural transmission (i.e. is not heritable). The two terms, *c_j_* and *g_j_* will be referred to as the culture-type and genotype, respectively. These terms only describe the state of the continuous variables, and are not meant to imply any particular mode of inheritance (e.g. haploidy, diploidy, etc.). Equation (1) is similar to the quantitative genetic formulation in Otto et al. (Otto et al., 1995). The culture- and geno-types are determined by the corresponding values in *j*’s genetic and cultural ancestors. We assume that a descendant’s culture-type and genotype are linear functions of her ancestors’ values given by

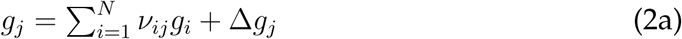

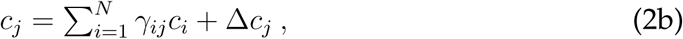

where *v_ij_*, *γ_ij_* ∈ [0, 1] and 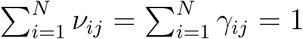; these values are the weights that describe the degree of influence an ancestor *i* has on descendant *j* in the genetic or cultural domain. When i is not a genetic ancestor to j, then *v_ij_ =* 0; when *i* is not a cultural ancestor, *γ_ij_* = 0. By normalizing these weights we have assumed that all individuals have at least one genetic and cultural ancestor. While this assumption is perfectly natural for genetic reproduction, one can imagine traits for which some individuals might receive no cultural input, or more cultural input than genetic. The delta terms, Δ*g_j_* and Δ*c_j_*, represent departures in *j* from the inherited genetic and cultural values. As an example, Δ*g_j_* may be nonzero in the event of mutation or recombination, while Δ*c_j_* may be nonzero due to individual learning or experience. This model generalizes that presented by El Mouden et al. (2014), though our analysis and conclusions differ.

Fitness captures the contribution of an ancestor to the next generation. In this model, that contribution, whether genetic or cultural, is determined by the weights given to an ancestor by her descendants (again, as in El Mouden et al. (2014)). Thus, the fitness of an individual in either domain of inheritance is simply the sum of the weights given to an ancestor by all descendants. Specifically, we define the genetic fitness of an ancestor *i* as 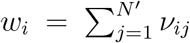 and the cultural fitness, 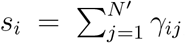, where the sums are taken over the descendant generation and *N*′ is the number of descendants. For example, for a haploid organism, all *v_ij_* are either 1 or 0, and *w_i_* is simply equal to the number of offspring; in the diploid, sexually reproducing case, *v_ij_ =* {0, 1/2}. For *s_i_,* the values will range from 0 to a maximum of *N*′, which occurs when *i* is the sole cultural ancestor of all descendants in the population. In the cultural domain, the definition of *s_i_* shows that the total amount of influence an ancestor i has on descendant phenotypes is what matters most, not just the number of individuals over which *i* has had some non-zero influence.

Using these definitions and equation 1, we can derive the following Price equation to describe the evolutionary change in the mean value of the phenotype (see SI-1),

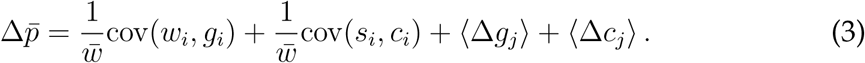

Note that angled brackets indicate averages over the descendant population, indexed by *j*. Just as in the standard Price equation, the covariance terms represent the effects of selection and drift (Rice, 2004) on evolutionary change. Importantly, we can separate the effects of differential reproduction, 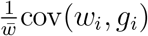, and differential influence in cultural transmission, 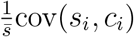. Note also that the mean fitness, 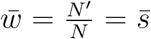, which is a direct result of the normalization conditions on *γ_ij_*⋅ and *v_ij_*, and again implies that everyone receives the same amount of cultural input as genetic input, and that cultural descendants must equal the mean number of genetic offspring. The remaining terms are the effects due to spontaneous departure from one’s inherited information, such as mutation or recombination in genes, or individual trial-and-error learning in culture. These terms differ somewhat from the transmission term in the standard Price equation, which is the fitness-weighted average departure of mean offspring phenotype from parental phenotype (*E*[*w*Δ*p*]). This difference results from the fact that we are measuring the differences (Δ*g_j_*,Δ*c_j_*) between an individual offspring’s type and it’s ancestral contribution.

The standard Price equation uses a single fitness measure and provides a mathematically valid description of evolutionary change. Why then do we need equation (3) that keeps track of two kinds of fitness? To see why, consider the population in Figure 1, which depicts an asexually reproducing population where descendants receive both genes and culture from ancestors, though not necessarily the same ancestors. The solid arrows indicate parent-offspring relationships, and the dashed arrows social learning relationships. Using only reproductive fitness (solid arrows) we could capture the evolutionary change with the standard Price equation: Δ*p̄* = *cov*(*w_i_*,*p_i_*) + *E*[*ω*Δ*p*] = –1/3 + 1 = 2/3. This expression allows us to see the effect of natural selection acting against the phenotype, but leads us to conclude that the transmission term, for reasons that are obscure, more than compensates for natural selection. Thus it appears that natural selection has been overtaken by a faulty inheritance system. However, computing the terms in equation (3), we have *cov*(*w_i_*,*g_i_*) = 0, *cov*(*s_i_*,*c_i_*) = 2/3, *E*[Δ*g*] = 0, *Ε*[*Δc*] *=* 0, and so *Δp̄ =* 0 + 2/3 + 0 + 0 = 2/3. Considering both genetic and cultural mappings,we see that there is in fact no natural selection on the phenotype in the genetic domain, and no flaws in either inheritance system; however, there is positive selection in the cultural domain that produces evolutionary change. This is a distinctly different cause than was revealed by considering only the reproductive fitness mapping. In summary, if the two modes of inheritance were not explicitly described as in equation (1), then a departure in phenotype from one’s genetic ancestors would include the effect of cultural inheritance, while a departure in phenotype from one’s cultural ancestors would include genetic inheritance. By explicitly accounting for both inheritance mechanisms, our approach avoids confounding their evolutionary effects.

We can use equation (3) to examine evolutionary change when there are conflicts between cultural and genetic selection forces. Is it possible for a trait that is favored by social learning but detrimental to reproductive fitness to evolve? For example, consider a socially acquired preference that leads to decreased reproduction, as in some cultural evolution models of the demographic transition (Ihara and Feldman, 2004; Kolk et al., 2014). Let higher values of *p* reduce fitness, that is to say, *cov*(*w_i_*,*p_i_*) *<* 0. Then we have the following condition,

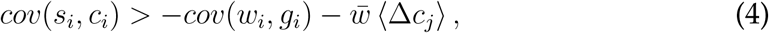

where we have ignored the genetic transmission term 〈Δg_j_〉 under the assumption that mutation and recombination effects are unbiased with respect to genotypic value. Putting aside for the moment the cultural transmission term, this condition states that the mean value of *p* can increase—despite reducing reproductive fitness—so long as the covariance between cultural value and influence on descendants exceeds the absolute value of the covariance between genotype and reproductive fitness. In essence, a loss in reproductive fitness can be compensated for by increased importance as a learning model. However, the cultural transmission term means that this condition will be harder to meet if social learning biases individuals toward lower cultural values than their learning models, for example, as a result of biased learning error (Henrich, 2004b).

Intuitively, whether individuals give higher or lower weights to ancestors with higher cultural values should determine the direction of evolution of *p*. This can be seen by observing that the cultural covariance term can be rewritten as (*see* SI-1)

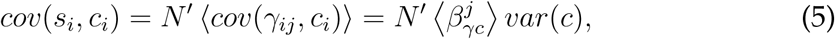

where again, brackets indicate the mean over the descendant population (index *j*) and *N′* is the descendant population size. The regression coefficient, 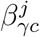 characterizes the learning rule employed by a descendant *j*; formally, it is the regression of the weight that descendant *j* ascribes to an ancestor on an ancestor’s cultural type. When this term is positive, it means that, on average, greater weight is given to ancestors with higher values of *c*. We can now rewrite eq. (4) as a new inequality that shows explicitly how strong the bias in favor of higher *c* must be in order for there to be positive evolutionary change,

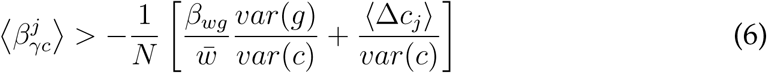

Condition (4) gives us the criterion for maladaptive phenotypes with respect to how ancestors’ *c* values translate into cultural fitness. The condition in (6) allows us to see the same condition from the ‘descendant’s point of view’. The population average of the learning rule employed by descendants determines the direction of evolutionary change. Importantly, we also see that the strength of the genetic selection term (first term inside the brackets) is modified by the relative variance in genotypes and culture-types. This is a result of having multiple selection terms in our Price equation. In fact, Hamilton (1975) pointed out a similar effect in his multilevel selection version of the Price equation, where the variances corresponded to individual and group level characters. One important difference is that while group and individual level variances are just different ways of partitioning the population variance (and hence have to add up to the total variance), here we have variances of two different variables whose values are unconstrained by one another.

### 2.1 Cultural Evolution of Altruism

Hamilton’s rule Hamilton (1964a,b) states that an altruistic allele will spread in the population when *rB > C,* where *B* is the fitness benefit to a recipient of altruism, *C* is the fitness cost to an altruist, and *r* measures the assortment between altruists (often interpreted as a relatedness coefficient). Cultural evolution theorists have claimed that altruism is more likely to evolve under cultural evolution, as this relatedness parameter for culture is likely to be higher than for genes (Fehr and Fischbacher, 2003; Boyd and Richerson, 2010; Henrich, 2004a). This claim implies that cultural evolution makes the spread of altruism possible even when the classical form of Hamilton’s rule does not hold (El Mouden et al., 2014), i.e. when genetic selection is opposed to altruism. To investigate this claim, the effect of evolutionary forces in the cultural and genetic domains must be compared directly, which has not been done before. In this section, we use our framework to derive the precise conditions under which cultural selection can favor altruism despite being opposed by genetic selection.

By altruism we mean here a behavior that reduces the fitness (genetic and/or cultural) of a focal individual while increasing the fitness of others, when the fitness effects of others on the focal individual are ignored (Hamilton, 1964a; Rousset, 2013). We assume that the fitness cost is both genetic and cultural. Let *p* now represent the level of altruistic behavior and the cultural and genetic fitnesses be given by the following equations:

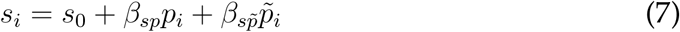

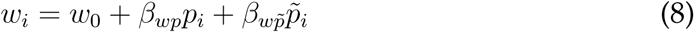

The tilde over a variable indicates the mean value of that variable across *i*’s neighbors. We have assumed both kinds of fitness are linear functions of an individuals own phenotype and the phenotypes of her neighbors, where *s_o_* and *w*_0_ are the baseline fitnesses. As in the standard derivation of Hamilton’s rule using the Price equation, it is customary to identify *β_wp_* and *β_wp̃_* as the cost (*C*) to an altruist and benefit (*B*) to recipients of altruism, respectively (Frank, 1998; Rice, 2004; McElreath and Boyd, 2008). We will use the same convention, but add subscripts to indicate costs and benefits to genetic *and* cultural fitnesses: *β_wp_* = – *C_g_*; *β_cp_ =* –*C_c_*; *β β_wp̃_* = *B_g_*; *β_cp̃_* = *B_c_*. By labeling these terms, we’ll be able to more clearly interpret our key results. We can derive the following condition (*see* SI-2),

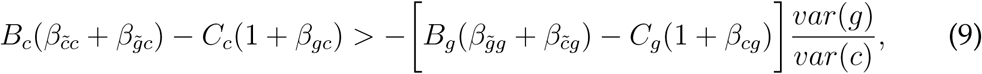

where we’ve ignored the transmission terms. Written this way, we can see that the left-hand side is the cultural selection coefficient, and the term in the brackets on the right-hand side is the genetic selection coefficient. One thing that is clear immediately is that the genetic selection coefficient is different than it would be under purely genetic transmission of the phenotype (i.e. *B_g_β β_g̃g_ – C_g_*, as follows from the canonical form of Hamilton’s rule). This is due to the presence of the additional regression coefficients *β_c̃g_* and *β_cg_*. The same can be said for the cultural selection coefficient, which would be *B_c_β_c̃c_* – *C_c_* under purely cultural transmission (El Mouden et al., 2014).

Taking first the LHS, we see three regression coefficients. The first, *β_c̃c_*, is the cultural relatedness term, and describes how likely actors are to behave altruistically toward individuals with similar culture-types. The second, *β*_*g̃c*_, is one of the gene-culture relatedness terms and captures the correlation between an actor’s culture-type and neighbor’s genotype. Thus, if individuals with higher culture-type values are more likely to direct their altruism towards those with higher genotypic values, the cultural fitness benefit is greater. The final regression coefficient, *β_gc_*, captures the correlation between an actor’s culture-type and its genotype. If altruism is costly both to cultural and genetic fitness, then a higher correlation between culture-type and genotype will make it even more difficult for altruism to evolve.

Now we turn to the genetic selection coefficient, given by the term in brackets on the RHS. The regression coefficient, *β_c̃g_*, is the regression of neighbor culture-type on focal genotype. The term *β_cg_* is the regression of focal culture-type and focal genotype. Both of these terms mean that the presence of cultural transmission changes genetic selection on altruism (i.e. genetic selection is no longer given by *Bgβ_g̃g_ – C_g_*), since there are now gene-culture relatedness terms to be taken into account. For example, if if individuals with an altruistic genotype are likelier to direct their altruism towards those with an altruistic culture-type then genetic selection can favor altruism even with low genetic relatedness (*β_g̃g_*).

The inequality states that the cultural selection coefficient must exceed the genetic selection coefficient scaled by the ratio of the variance in genotypes to cultural types. Thus, even relatively weak cultural selection can overcome genetic selection if the variance in culture-types is sufficiently high compared to the variance in genotypes. Below we will explore the consequences of (9) using three simple illustrative models.

## 3 Illustrative models

We imagine a population of haploid individuals interacting assortatively in each generation. These interactions determine the reproductive output of each individual and, potentially, their cultural influence on the next generation. Each individual possesses two loci with a single ‘allele’ at each locus. At the first locus, alleles are transmitted genetically, from a single parent to her offspring; at the other locus, a ‘cultural allele’ is acquired from a single cultural parent. An individual’s phenotype is determined by the combined additive effect of the alleles at the two loci in the following way: when two individuals interact they play a prisoner’s dilemma; each individual employs a mixed strategy where the phenotype, *p*, is the probability of playing ‘cooperate’. Those with both the genetic and cultural alleles for altruism play a pure strategy of cooperate; those with only the genetic or cultural allele, play cooperate half of the time; finally, an individual that lacks both the genetic and cultural alleles will play a pure strategy of defect. Thus we have four types of individuals in the population {0, 0}, {0, 1}, {1, 0}, {1, 1}, with phenotypes *p*_00_ *=* 1, *p*_01_ *= p*_10_ *=* 1/2, *p*_11_ *=* 1.

An individual of type *ψ* has an expected reproductive fitness of

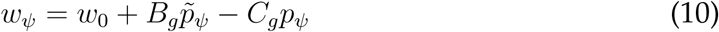

where *w*_0_ is the baseline fitness, *p̃_ψ_* is the expected phenotype of a type *ψ* individual’s opponent in the game, and *p_ψ_* is the phenotype of a type *ψ* individual.

Players in the model interact assortatively with respect to both genes and culture. The probability that a player encounters an opponent of the same genotype is *f*_g_, while the probability for culture-types is *f_c_*. If individuals were interacting with kin, *f_g_* would be the probability of being identical-by-descent, and *f_c_* would be the analogous value computed for a cultural genealogy (Aguilar and Ghirlanda, 2015). We allow for these assortment parameters to vary by domain because population structure may vary with respect to the two systems of inheritance. For example, in a genetically panmictic population in which cultural learning only takes place within demes, *f_g_* is 0 while *f_c_* is nonzero. These two values allow us to consider a broad range of population structure models. We assume that with some probability, *f*_g_, an individual chooses a partner of identical genotype, and otherwise selects her partner at random (with an analogous situation for culture-type). We then compute the probability of having a player of a certain type will be conditional on one’s own type. Thus, the probability that a type {1, 1} interacts with another {1, 1} is,

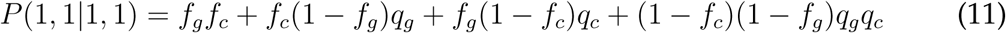

where *q_g_* and *q_c_* are the population frequencies of the genetic and cultural altruistic alleles. The first term is the probability that two {1, 1} individuals are identical due to assortment; the second is the probability of being identical due to assortment for culture but not genes; the third is the probability of being identical due to assortment for genes and not culture; and the final term is the probability of not being identical due to assortment either genetically or culturally. These conditional probabilities then determine the expected phenotype of an individual’s opponent in the game, *p̃_ψ_*. Further details on the model are provided in SI–3.

Offspring inherit their parent’s genetic allele. They must then choose a cultural model whose allele they will inherit at the cultural locus. In the next two subsections, we consider two models differing in how cultural parents are chosen. In both, cultural models are independent of the genotype of the offspring. In section 4, we consider the more general case where there is genetically biased cultural transmission.

### 3.1 Model 1: Neutral cultural trait

First, we assume that the cultural propensity for altruism is neutral with respect to cultural fitness. In other words, ancestors are chosen as cultural parents without regard to their cultural traits, so the probability of acquiring the cultural propensity for altruism is *q_c_,* the population frequency of the cultural allele in the parental generation. We can use (9) to determine the condition for the increase in the altruistic phenotype by multiplying both sides of the inequality by *var*(*c*) and computing the covariances directly from the model. We have no cultural selection, so *B_c_ = C_c_* = 0. Since culture is chosen at random, genetic and cultural type are uncorrelated, so that *cov*(*c*, *g*) = *cov*(*c̃*, *g*) *= cov*(*g̃, g*) = 0. Thus, (9) reduces to

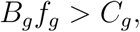

the canonical form of Hamilton’s rule. This result follows directly from the cultural allele being chosen at random. Under random copying the expected change in the frequency of the cultural allele is zero and the only change in mean phenotype will be due to changes in the frequency of the genetic allele. Further, with no correlations between the genetic and cultural allele, the only forces affecting the evolution of the genetic allele will be the reproductive fitness effects. However, it should be noted that due to the dual inheritance of altruism, the value of the phenotype may be maintained at significant levels in the population if the frequency of the cultural allele is high. Take the extreme case where *q_c_* =1. Even if the inequality above is not met and the genetic allele is driven to extinction, the cultural allele will be unaffected and the mean value of the phenotype in the population will be *p̄= q_c_*/2 = 1/2. In other words, there will be no perfect altruists, but everyone will be a ‘half’ altruist. As the mean reproductive fitness, *w̄*, depends on the mean phenotype, this could have important implications for population growth, including eventual extinction.

### 3.2 Model 2: Cultural prisoner’s dilemma

Next, we consider a case where offspring no longer choose their cultural parent at random. In particular, we assume that individuals meet to play the prisoner’s dilemma, this time with respect to both reproduction and cultural propagation. For simplicity, we’ll imagine individuals producing cultural ‘gametes’ or behavioral tokens that can then be acquired by offspring. The number of tokens an individual produces will affect her probability of being copied. This can be interpreted as her visibility, or salience as a cultural model. The expected number of cultural gametes, *z*, that an individual of type *ψ* produces is,

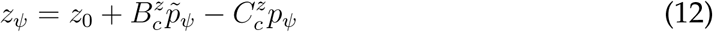

The terms 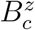 and 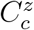 are the gametic fitness benefit and cost (i.e. the regression of the number of cultural gametes produced on neighbor phenotype and focal phenotype, respectively), with 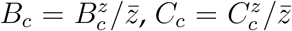 (see SI-4). Recall that in the previous section cultural fitness was defined in terms of the total influence 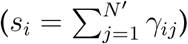 an ancestor has on the descendant population. In this model, offspring have a single cultural ancestor (i.e. *γ_ij_ =* 1), and *s_i_* is just the total number of descendant individuals who count i as an ancestor. The number of offspring available as cultural descendants is determined by the reproductive output of the population, thus,

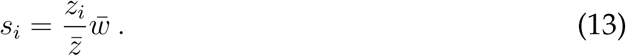

Substituting (12) and (13) into the gene-culture Price equation and making simplifications (*see* SI-4 for details), we obtain:

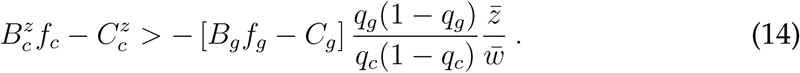

We now have an additional scaling term, *w̅/z̅* = *s_i_*/*z_i_*, the number of cultural descendants per cultural gamete produced. This term converts the payoff in the cultural Prisoner’s Dilemma game (*z_i_*) to the true cultural fitness (*s_i_*), which in this case is the number of individuals that count *i* as an ancestor.

Above, we have written out the variance terms explicitly. For given cultural and genetic selection coefficients, the ratio of the variances in (14) means that the effect of cultural selection will be maximized when the genetic allele is at very high or very low frequency (*q_g_* close to 0 or 1) and *q_c_ =* .5.

In Figure 2, we plot the the values of the total selection coefficient on the altruistic phenotype (i.e. LHS *minus* RHS in (14)) against the frequency of the genetic allele, *q*_g_. The frequency of the cultural altruistic allele is set to .5 to maximize *var*(*c*), which allows us to restrict the values of *var*(*g*)*/var*(*c*) between 0 and 1. As *var*(*g*) is minimized at *q_g_* = 0, 1, selection is convex in *q*_g_. This is generally true so long as the cultural selection coefficient is positive and the genetic selection coefficient is negative. Thus, when *q_c_* = .5, cultural selection will have the strongest effect when the genetic allele is extinct or at fixation.

**Figure 2:**
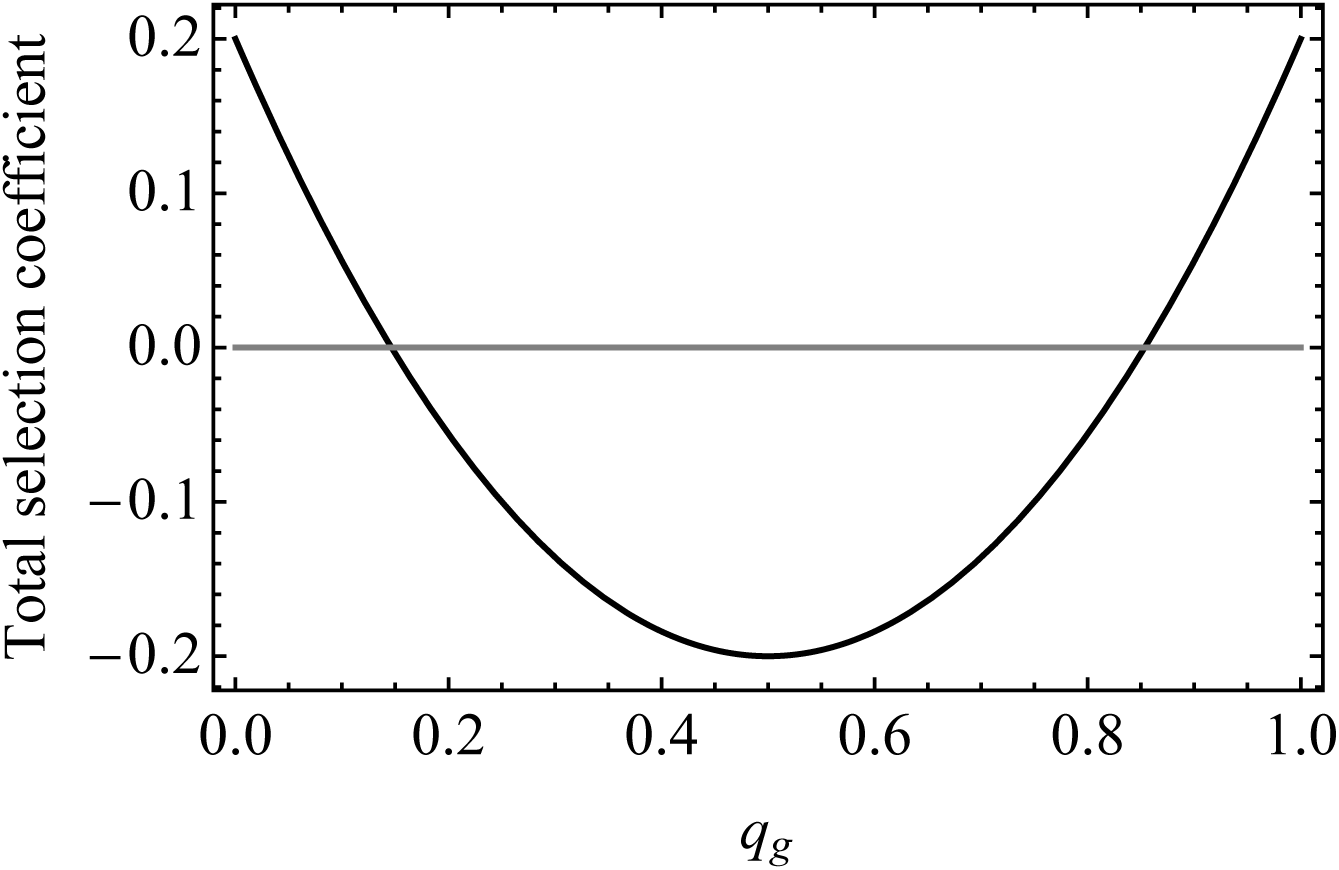
Total selection coefficient (LHS-RHS in (14)) for 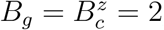, 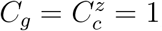, *r_g_* = .3, *r_c_* = .6, and *q_c_* = .5, to maximize *var*(*c*). Total selection coefficient is maximized when *q_g_* is at its minimum or maximum values. Parameter changes adjust the range, but not the shape of the function, so long as 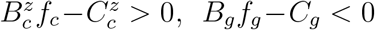, *B_g_f_g_*−*C_g_* < 0 (i.e. there are some parameter combinations for which all values are negative/positive but the curve remains convex).

We defined altruism with respect to both cultural and genetic fitnesses. In model 1, cultural transmission was neutral with respect to the altruistic phenotype, while in model 2 there was also a direct cultural fitness cost to the phenotype. It is possible that a phenotype may be beneficial in the cultural domain while detrimental to reproduction. In this case, we simply change the sign of the cost term on the LHS of (14) and see that it makes the condition easier to meet. Conversely, we could imagine a behavior that is costly with respect to cultural fitness and entirely beneficial in the genetic domain, which would again make condition (14) more difficult to meet. This highlights the importance of specifying the fitness consequences of a trait in each domain of inheritance.

## 4 Gene-Culture covariance

In the two illustrative models above, we assumed that genetic and cultural transmission were independent, i.e. having a particular genetic type had no effect on the probability of acquiring a particular cultural type. This allowed us to ignore the covariance between genetic and cultural types. Now we allow for genes and culture to be nonindependent. In particular, we assume that with probability k a newborn individual will acquire the cultural allele that mirrors her genetic allele. For example, an individual born with the genetic allele for altruism (*g* = 1), will acquire the cultural allele for altruism with probability *k*. With probability 1 – *k*, she acquires her cultural allele randomly from the cultural gamete pool. This kind of non-random learning can result, for example, from a genetic predisposition towards particular cultural models, such as when the locus that determines cultural learning is linked to the locus that determines the altruistic genotype. The effect of this construction is to introduce a correlation between genotype and cultural type.

The full recursion equations are given in a Mathematica notebook in the SI. In SI-5 we show that the altruistic phenotype increases when:

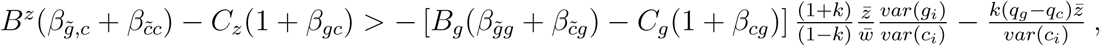

which evaluates to:

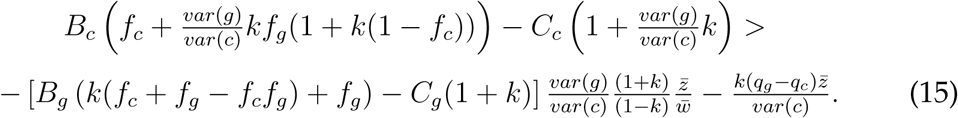

When *k* = 0 the above expression reduces to (14). When *k* = 1, the RHS goes to positive or negative infinity, depending on the sign of genetic selection; when genetic selection is negative, the condition becomes impossible to meet. Thus, the effect of k, the correlation between genotype and cultural type, is to make it more difficult for cultural selection to overcome natural selection when they are in conflict. This is intuitive as higher *k* makes the cultural inheritance system more coupled to the genetic inheritance system.

As in the general condition, (9), the genetic selection term (i.e. the term on the RHS in brackets) is different from what it would be under purely genetic transmission of altruism. One way to see this effect is to fix *B_g_* and *C_g_* and see above what value of *f_g_* the genetic selection coefficient is positive, which we’ll call 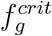. For example, under purely genetic transmission of altruism, for *B_g_* = 2, *C_g_* = 1, then 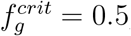; for all *f_g_ >* 0.5 genetic selection favors altruism. In figure 3 we plot 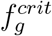 for the same values of *B_g_* and *C_g_* and varying *r_c_* and *k*. For all constellations of *f_c_* and *k* we see that 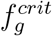 under gene-culture transmission is always lower than 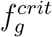 under purely genetic transmission (in fact, in can be shown that for *B_g_ > C_g_*, the critical value will always be lower). Thus, cultural transmission affects the evolution of altruism in two important ways: (1) by introducing a cultural selection force that may overcome genetic selection; (2) by changing the nature of genetic selection itself. This latter effect means that for given values of *B_g_* and *C*_g_, genetic selection may be positive in the presence of joint cultural and genetic transmission when it would have been negative under purely genetic transmission.

**Figure 3:**
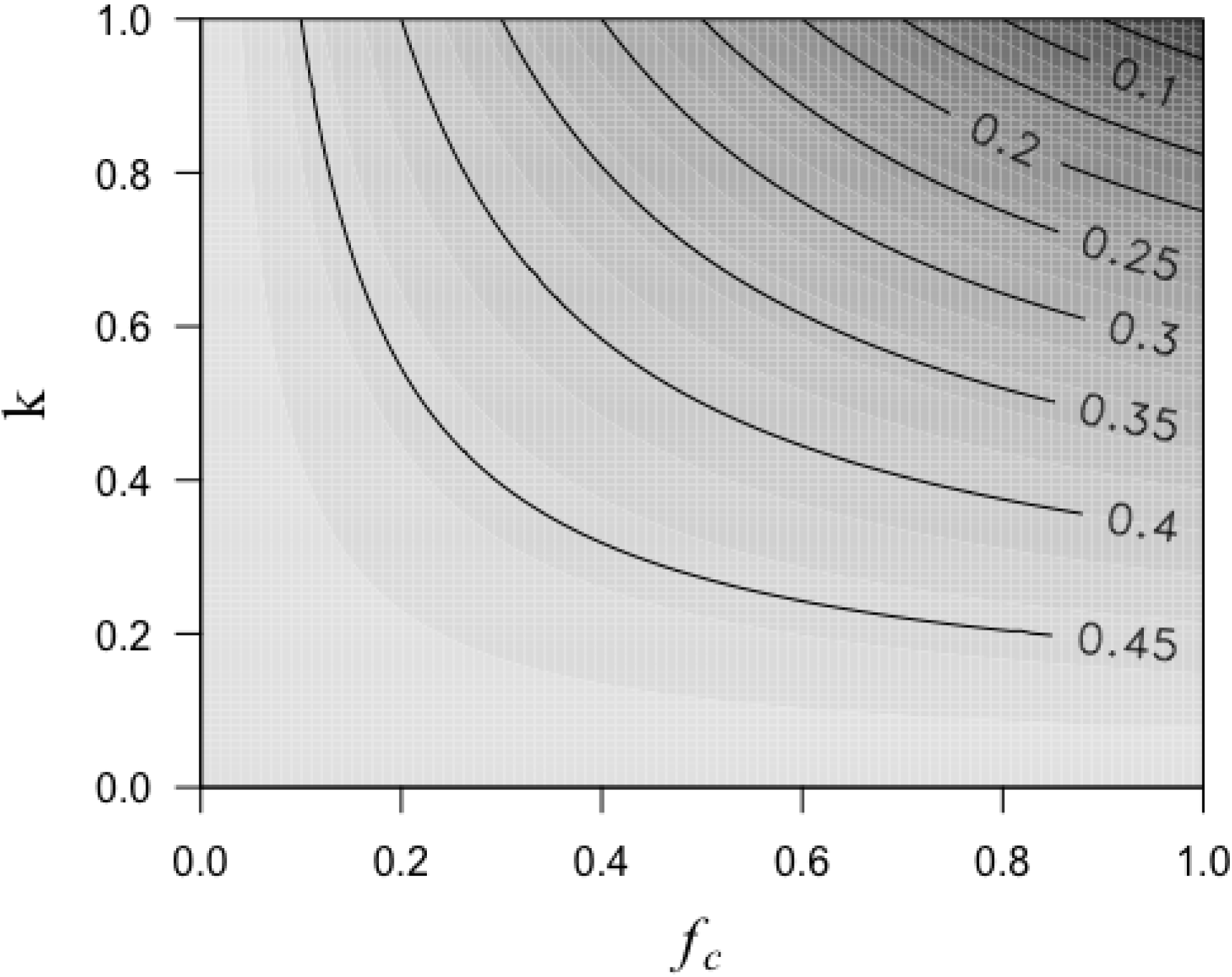
Critical values of 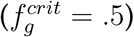 for *B_g_* = 2, *C_g_* = 1 and varying *r_c_* and *k.* For all combinations of *r_c_* and *k*, 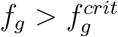 is lower than for what it would be under purely genetic transmission of altruism 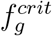. Values 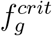 mean positive genetic selection.

## 5 Non-additive phenotypes

The results described above all assumed an additive phenotype function, which is a standard starting point in social evolution and population genetics theory (Van Cleve, 2015). However, biological reality may be much more complicated, particularly when trying to incorporate the effects of multiple inheritance systems. One way to deal with this problem in evolutionary theory has been to observe that most genetic variants have small effects on phenotypes and genetic variation in the population is small, in which case, an additive approximation gives satisfactory results (Taylor and Frank, 1996; Akçay and Van Cleve, 2012). In this section, we translate this approach to phenotypes that are jointly determined by genes and culture.

We begin by assuming that an individual descendant *j*’s phenotype is given by a function *p_j_* (*c_j_*, *g_j_*), where the arguments are a descendant’s heritable cultural and genetic information. This information in turn is a function of the heritable cultural and genetic information of that individual’s ancestors, which implies that we can instead write the phenotype mapping function as *p_j_* (*c*_1_,*…,c_N_,g*_1_,*…, g_N_*), a direct function of the ancestral culture-types and genotypes. Assuming that all *p_j_* are differentiable with respect to ancestral values, and that the variances in *c* and *g* are small, we can make a first-order Taylor approximation of *p_j_* around the point (c̄,ḡ)=(*c̄,…,c̄,ḡ…,ḡ*). We then substitute this expansion into 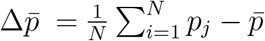 to arrive at a Price equation for the non-additive case (*see* 5),

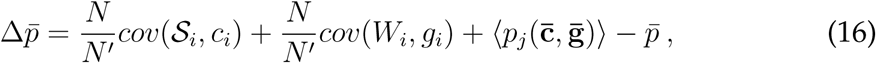

where 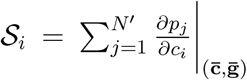 and 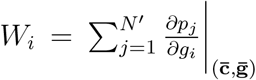, refer to *generalized fitnesses* in the sense that we are measuring not only the number of descendant individuals an ancestor has, but also the combined effect of that ancestor on her descendants’ phenotypes. For example, in a haploid genetic model in the absence of mutation, where the ‘phenotype’ of interest is just the genotype, then 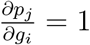 when *i* is a genetic ancestor of *j*, while 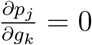 for all individuals *k* that are not genetic ancestors to *j*. In this case, the generalized fitness just reduces to the number of descendant individuals who count i as an ancestor. Similarly, in the model presented in the first section, the partial derivative of the phenotype function *p_j_* with respect to *c_i_* will yield *γ_ij_*, and *S_i_* = *s_i_.* The advantage of this formulation is that more complicated phenotype mapping functions can be incorporated into the idea of a generalized fitness. The formulation also allows for the generalized fitness to be negative, as might happen in a case where ‘offspring’ react by contrasting themselves with parental individuals (Findlay, 1992). This generalized fitness concept captures the causal effect of an ancestor on a descendant individual’s phenotype, where it induces a positive or negative correlation in their phenotypes, a feature missing from more restrictive fitness concepts.

Equation (16) looks similar to equation (3); first, we have two covariance terms that account for the effect of selection (now with respect to generalized fitness). We’ve replaced the inverse of the mean fitness with a more direct measure of population growth, *N/N′*; this is because generalized fitness refers to the effect of an ancestor on the phenotypes in the next generation, and is no longer synonymous merely with her contribution to the growth of the population. The remaining term, 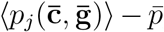, is the difference between (1) the average phenotype that would occur if every individual inherited the mean values of *c* and *g*, and (2) the mean phenotype among ancestors (*p̄*). This isolates the effect of the phenotype functions among descendants, *p_j_*, on evolutionary change and can be seen as analogous to the traditional transmission term in that it captures the effect of constructing phenotypes from inherited information on evolutionary change.

From eq. 16 we can simply derive a condition for the evolution of a maladaptive trait. When Δ*p̄* > 0, we have,

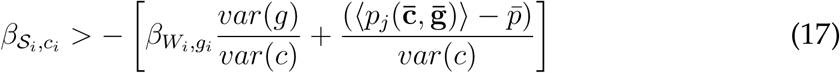

This result is exactly analogous to (4) in the first section and can be summarized similarly: a loss in generalized reproductive fitness can be compensated for by a gain in generalized cultural fitness. Again, we have assumed that the rules of transmission remain constant over the timescale being considered in the Price equation.

This approach could of course be extended to higher order expansions of the phenotype function: in Section 5 we show that the infinite expansion of the phenotype function leads to a more precise definition of generalized fitness than the approximation given in this example. Most importantly, without making assumptions about either the phenotype mapping function or the fitness function, we have shown an important relationship between these two fundamental concepts in evolutionary theory.

This approach bears some resemblance to the model presented by Richerson and Boyd in (Richerson and Boyd, 1978), where generic phenotype functions depended on genotype, cultural-type, and a third argument called the penetrance, which determined the relative influence of either inheritance system on phenotype. In the foregoing presentation this penetrance value is implicit in the phenotype function.

## 6 Discussion

### 6.1 Co-inherited Behaviors

Our model was inspired by the fact that many complex behavioral traits, from psychological traits, to disease risk, are likely affected by both genetic and cultural transmission. For instance Levy et al. (2016) found that culturally transmitted negative stereotypes about aging predicted later biomarkers of Alzheimer’s disease (reduced hippocampal volume and increased amyloid plaque development). Our model indicates that if these negative stereotypes are culturally selected for, perhaps because a rapidly changing environment makes older individuals seem less valuable as cultural role models, then cultural selection can increase the incidence and severity of Alzheimer’s biomarkers, especially as natural selection is likely to be very weak in this case. In another interesting example, Kong et al. recently showed evidence for selection against genetic variants associated with educational attainment, a complex trait which is clearly influenced by social learning (Kong et al., 2017). If more highly educated individuals exercise greater influence on the educational attainment of others then cultural and genetic selection will be in direct conflict. Our model suggests that measuring the relative variance in culture and genes for this co-inherited trait (educational attainment) is an important next step in understanding its evolution.

Research into the evolutionary basis of human behavior has long puzzled over the existence of maladaptive behaviors (Glanville, 1987; Logan and Qirko, 1996). These are behaviors that persist via cultural transmission despite detrimental reproductive fitness effects, such as clubbing pregnant women to induce birth in Colombia (Reichel-Dolmatoff and Reichel-Dolmatoff, 2013), unhygenic neonatal care practices in Bangladesh (McConville, 1988), and folk medical practices like ingesting rhino horn (Ayling, 2013) or bloodletting (Wootton, 2007). While these practices are likely spread almost exclusively by cultural transmission, they may be influenced by genetic inheritance via broader behavioral traits with a significant genetic component, such as risk-taking, which also shows cross-cultural variation (Weber and Hsee, 1998; Hsee and Weber, 1999). Fertility behavior provides another potential example of a dually inherited trait. The reduction in fertility seen across the globe, known as the demographic transition, may be a result of cultural selection overcoming genetic selection for higher fertility. Our model demonstrates more broadly the possibility that maladaptive behavioral traits may evolve under dual transmission, despite their reproductive fitness costs. In fact, using our Taylor expansion approach for non-additive phenotypes, the Kolk et al. model can be expressed in terms of our Price equation.

### 6.2 Relative strength of genetic and cultural selection

Explicitly modeling genetic and cultural inheritance for a co-inherited trait did not only reveal the effect of each inheritance system on evolutionary change; it also revealed the effect of each inheritance system on the other. As we saw in sections 2.1 and 4, the presence of cultural transmission changed the genetic selection coefficient from what it would be under purely genetic transmission (the same is true with respect to purely cultural transmission). The addition of another inheritance system for a single trait meant that there was both an additional selection force to be considered, and that genetic selection itself took a different form that could favor the evolution of a trait when purely genetic transmission would not.

Our results show the importance of the ratio of genetic to cultural variance in scaling the effect of genetic selection. It is interesting to consider empirical estimates of cultural and genetic diversity to gauge the expected relative strength of genetic selection. Bell et al. compared *F_st_* values for culture and genes in populations using the World Values Survey (Bell et al., 2009). Their results suggested greater-between population variation in culture than in genes. Unfortunately, these results say little about the within-group variance in culture relative to genes. Other studies have shown parallels in the patterns of linguistic and genetic diversity (Perreault and Mathew, 2012; Longobardi et al., 2015; Creanza et al., 2015; Hunley et al., 2008), but again provided no information about the ratio of genetic to cultural variance. However, this question is well-suited to empirical study; given our results, empirical estimates of the ratio can shed light on qualitative expectations about the evolution of behavioral traits.

The ratio of genetic to cultural variance also has an important relationship to the narrow-sense heritability (*h*^2^), which measures the proportion of phenotypic variance attributable to the ‘heritable’ component of phenotype (Falconer and Mackay, 1996). In a series of papers, Danchin and co-authors (Danchin and Wagner, 2010; Danchin et al., 2011, 2013) introduced the idea of ‘inclusive heritability’, which partitions the variance in the heritable component of phenotype into the contributions from each system of inheritance. This allows for narrow-sense heritability to be expressed as the sum of the heritabilities in each domain (assuming no interactions between the inheritance systems). In our model, this means 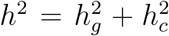 (where 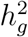 and 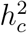 are the genetic and cultural heritabilities). The ratio of these heritabilities is exactly the term that appears in our results as the scaling factor of genetic selection, further demonstrating the importance of inclusive heritability when considering evolutionary outcomes.

### 6.3 Related results

Nearly forty years ago Richerson & Boyd showed that under certain conditions, the equilibrium value of a trait that is both genetically and culturally inherited could be that which optimizes cultural fitness (Richerson and Boyd, 1978). But surprisingly, given the intense attention gene-culture coevolution received, very little theoretical work has been done to follow up on the evolution of phenotypes that are directly co-inherited, as opposed to culturally inherited behavioral phenotypes and genetically inherited learning rules (e.g., as in Lehmann and Feldman, 2008). Findlay (Findlay, 1992) modeled gene-culture transmission of a phenotype in a structured population. He found that when individual level selection is weak and migration low, gene-culture transmission was more favorable to the evolution of altruism than genetic transmission alone. This is because the effect of migration is to erode between group variance. In our paper the total population variance in culture was shown to have an effect on making the evolution of an altruistic phenotype easier, and we implicitly included the effect of population structure in our illustrative models. Another paper close to our model is that of Lehmann et al. (2008), who model the evolution of a purely culturally inherited altruistic behavior in a subdivided population. In their model, Lehmann et al. consider the phenotype to be affecting either only cultural or reproductive fitness, and assume no genetic contribution to the phenotype. As such, their model can be recovered by modifying our Price equation (9) (by setting *p_i_* = *c_i_*, which replaces the right-hand side of the condition with zero). Importantly, biological offspring in Lehmann et al’s model serve as “vectors” of the cultural types of their parents, which means that even when their cultural trait only affects reproductive fitness, our corresponding *B_c_* and *C_c_* terms would be non-zero. That means that the transmission rate of different cultural types are not the same over the entire lifecycle, and there is cultural (but no genetic) selection.

More recently, El Mouden et al. used a Price equation to describe cultural evolution and made some claims that at face value differ from our results. In particular, they claim that cultural selection can only increase genetic fitness (e.g., through altruism that benefits others) if cultural and genetic fitness are tightly coupled. We have instead shown that such co-inherited altruism can increase through cultural selection even when opposed by genetic selection. The source of the discrepancy lies in the fact that El Mouden et al. base their statements on the *partial change in genetic fitness due to selection* (cf. their equation S12), which by Fisher’s fundamental theorem (Fisher, 1930; Bijma, 2010) is always positive, even though total mean fitness can be increasing or decreasing. To give the simplest possible example, consider our equation (14), and an altruistic trait with *B_g_*>*C_g_* but *f_g_B_g_* < *C*_g_. This trait would be opposed by genetic selection, but if it spreads through cultural selection (which happens when 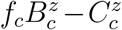 is sufficiently large) it would increase the mean genetic fitness, despite the fact cultural and genetic fitness are negatively correlated. This is a direct consequence of the well-known fact the total change in fitness includes changes in the population composition (e.g., social effects) that are ignored in the Fisherian partial change. This bears directly on the claim of El Mouden et al. (on p. 235) that cultural selection opposed by genetic selection cannot explain the success and spread of human populations through increased reproductive fitness, because the latter is clearly an argument about realized (or total) fitness, and not about partial changes. That claim is false as the above argument illustrates: cultural altruism, even if it is opposed by genetic selection *can* increase the mean fitness of populations. Furthermore, because El Mouden et al. do not consider co-inheritance of a phenotype underlying fitness along with potential gene-culture covariances, they overlook the possibility that cultural transmission and assortment might alter the direction of genetic selection. Thus, contrary to the arguments of El Mouden et al., the sovereignty of genetic selection over cultural selection is far from absolute, and a careful accounting of the operation of both modes of transmission is needed.

### 6.4 Limitations and extensions

The additive model we used in this paper is both the simplest model and a natural extension of the standard assumption in quantitative genetics (Falconer and Mackay, 1996). However, even under this simple model we observed some nontrivial results. In the general model we assumed that mean cultural fitness was equal to the mean reproductive fitness (i.e. *s̄ = w̄*). This was convenient and simply resulted in every individual receiving some cultural input, though there exist traits for which some individuals may never receive cultural input. For example, though underlying genetic variation may determine one’s reading ability, one may never be taught to read. In these cases, the equality of *s̄* and *w̄* will not necessarily hold. Our general framework could easily be extended to compass this possibility. Indeed, we discarded this assumption in our second illustrative model; when the mean number of replications for culture and reproduction were not the same—in this case *z̄* and *w̄*—the conversion factor *z̄*/ *w̄* scaled the effect of genetic selection. In the event that cultural replication might affect fewer individuals than are actually born, *z̄/w̄* < 1, and the effect genetic selection is further reduced.

In the course of deriving our results on the effects of selection, we often ignored the transmission terms, 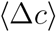 and 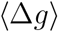. In relatively simple genetic systems, it may be safe to assume that the expected difference between parents and offspring is zero. However, culture very often makes this assumption untenable, as the cultural transmission system allows for biased or directed ‘mutation’ in the form of individual learning and other factors. For example, individuals may systematically differ from their parents because they learn more appropriate responses to their environment through their own trial-and-error learning. El Mouden et al. (2014) offered an interpretation of the transmission term as evolved biases in favor of reproductive-fitness-maximizing behaviors. However, biases (or individual learning) may not always produce reproductive fitness-maximizing biases, especially under frequency dependence. Meanwhile, Henrich (2004) took the transmission term to represent systematic error in cultural learning that biased individuals to trait values lower than their cultural parents. These examples hint at the diverse interpretations that can be ascribed to the transmission term, particularly in lieu of empirical evidence on how a specific trait is passed on. These effects also present important future directions for a more complete framework of gene-culture co-evolution.

In our section on non-additivity, we took an unusual approach to deriving the Price equation. Most models of social evolution make an explicit assumption about the fitness function (e.g. linearity, as in our derivation of the gene-culture Hamilton’s rule) and an implicit assumption about the phenotype function (e.g. *p =* g, as in the phenotypic gambit). By contrast, we made no assumptions about the form of the phenotype function, with the exception of infinite differentiability, and were able to derive a definition of fitness that similarly relied on no previous assumptions about the fitness function. This approach demonstrates the relationship between how phenotypes are actually constructed from inherited information and fitness itself. Also, our notion of generalized fitness captures both the effect of reproduction (the number of individuals who receive any heritable information from an ancestor) and the effect of transmission (how much heritable information flows from an ancestor to a descendant). The relationship between generalized fitness and other important fitness concepts, such as inclusive fitness, is worth exploring, but beyond the scope of the present paper.

Lastly, we took the transmission rules for both genes and culture to be stable over the timescale assumed in the Price equation. While this is the standard assumption for genetic systems, the long-term evolution of culture will be determined by the ways in which individuals acquire cultural information, a trait that may itself be culturally or genetically transmitted. Exploring the evolution of transmission rules in the context of a trait that is co-inherited is an important future direction for this work.

### 6.5 Conclusion

The Price equation offers a general statement of how evolutionary change can be partitioned among different evolutionary factors (Frank, 2012). We derived a Price equation for the evolution of a trait that is transmitted via both modes of inheritance. In using the Price equation, we have offered a general framework for partitioning the evolution of a co-inherited trait across the distinct causes (selection, transmission effects, etc.) in each domain of inheritance. Given the evidence for the long evolutionary history of cultural transmission in the human lineage (Lind et al., 2013), it is likely many behavioral traits evolved under the combined influence of genetic and cultural transmission. As the importance of non-genetic inheritance systems becomes clearer, we propose that accounting for multiple inheritance systems explicitly, as we do in our framework, will contribute to a better theoretical understanding of the evolution of these traits.

## Supplementary Information

### SI-1 Derivation of Gene Culture Price equation

The phenotype of individual *j* is given by,

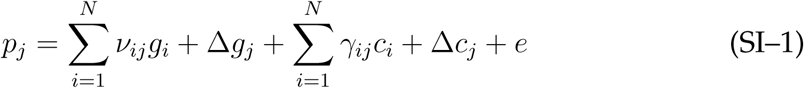

where the coefficients *v_ij_* and *γ_ij_* represent the influence an ancestor *i* has on descendant *j* in the genetic and cultural domains, respectively (Note: 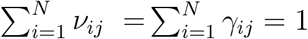)^1^. The mean value of *p* in the descendant generation is,

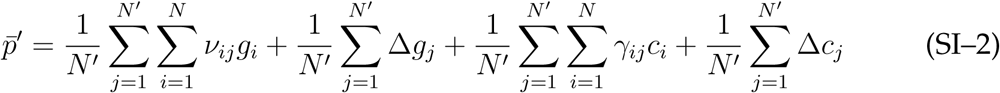

where *e* is assumed to have mean zero. Reversing the orders of the double sum terms and noting that 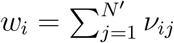, and 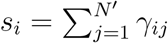, we can rewrite eq. SI-2 as,

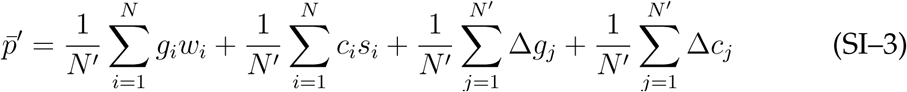

Using the definition of covariance (*cov*(*x*, *y*) *= E*[*xy*] *– E*[*x*]*E*[*y*]) we can replace the first two terms on the RHS,

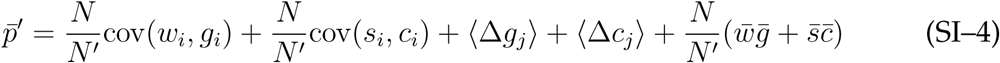

The angle brackets here mean averages over the descendant population. Noting *Nw̄ = Ns̄ = N^′^* we can rewrite the final term on the RHS as *ḡ* + *c̄*. subtracting the mean phenotype in the ancestral population, *p̄* = *ḡ* + *c̄*, we have (3).

The cultural covariance term in (3) takes the ‘ancestral’ point of view, in that it includes ancestral cultural values and their fitnesses. However, we can be re-express this term from the descendant point of view with the following quick restatement (note: overbars are ancestor averages and brackets are descendant averages),

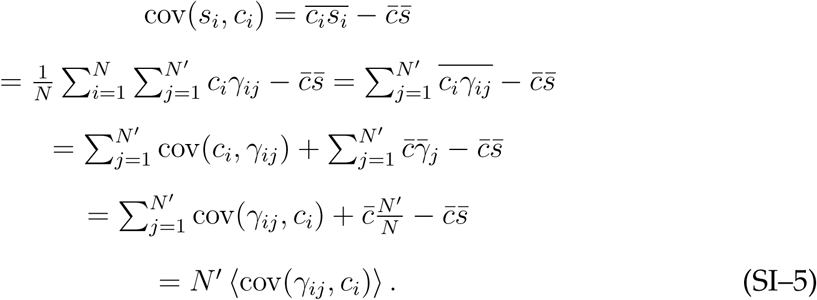

For clarity, the final mean of the covariance term is taken over the descendant population.

### SI-2 Derivation of Gene-culture Hamilton’s rule

We begin with the following cultural and genetic fitness functions:

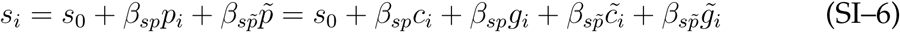

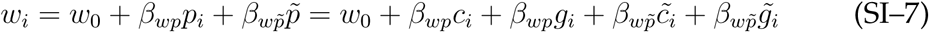

The tilde over a variable indicates the mean value of that variable across *i*’s neighbors. We have assumed both kinds of fitness are linear functions of an individuals own phenotype and the phenotypes of her neighbors. As in the standard derivation of Hamilton’s rule using the Price equation, it is customary to identify *β_wp_* and *β_wp̃_* as the cost (*C*) to an altruist and benefit (*B*) to recipients of altruism, respectively. We will use the same convention, but add subscripts to indicate costs and benefits to genetic *and* cultural fitnesses,

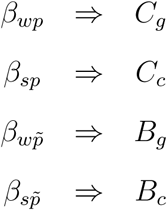

Substituting SI-7 into our Price equation in 3, and ignoring the transmission terms, we have,

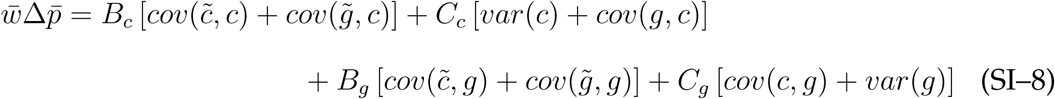

The equation above allows us to derive a condition for the evolution of the altruistic trait p in the population. Using *cov*(*x*,*y*) *= β_xy_var*(*y*), where *β_xy_* is the linear regression coefficient of *x* on *y*, and dividing through by *var*(*c*), we can rearrange the above expression to find,

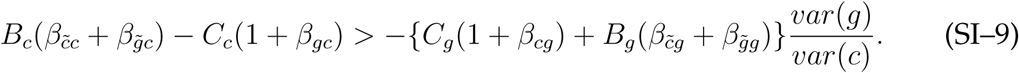

### SI-3 Model 1

We imagine a population of haploid individuals who, once born, select a cultural parent to copy. Each individual has two loci with a single allele present at each. The allele at the first locus is genetically transmitted while the allele at the second is received from a cultural parent. Individuals interact assortatively, with some probability of being genetically identical due to assortment, (*f*_g_), and culturally identical due to assortment, (*f_c_*). At discrete time steps individuals meet a random kin member and play a prisoner’s dilemma according to a mixed strategy. The phenotype, *p*, is the probability of playing cooperate. The two loci mean four types of individuals {0, 0}, {0, 1}, {1, 0}, {1, 1}, with phenotypes, *p*_00_ *=* 0, *p*_01_ = 1/2, *p*_10_ = 1/2, *p*_11_ = 1.

The expected reproductive fitnesses for each type are

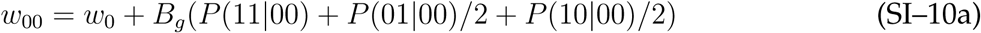

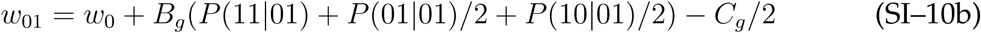

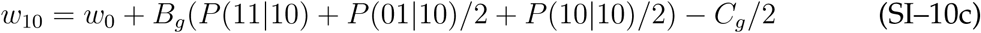

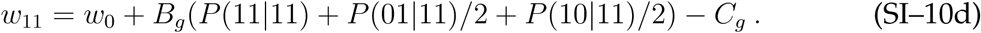

The conditional probabilities are probability of encountering a certain type given one’s own type. For example, *P*(10|00) denotes the “probability of encountering a {1, 0} given that the player is a {0, 0}.” Rather than enumerate all of these conditional probabilities we take advantage of the following identity:

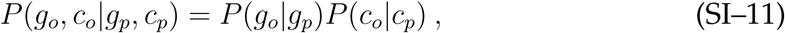

where the *o* subscript indicates the opponent and *p* the player. We need only specify the following conditional probabilities,

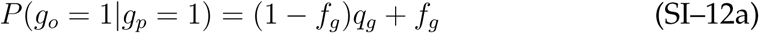

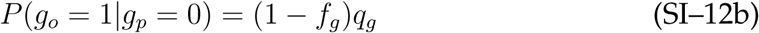

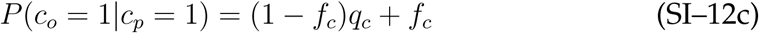

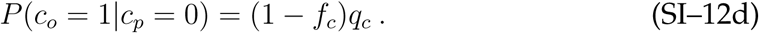

Note that the remaining marginal conditional probabilities are given by

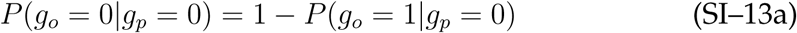

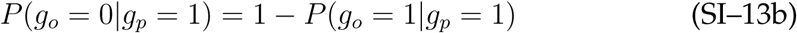

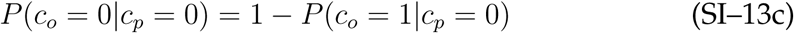

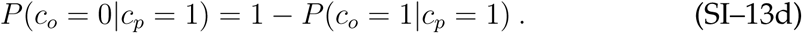

Using (SI–11) we can calculate all the conditional probabilities of encounters between types.

For each type we can write the following recursions for the frequency at the successive time step by simply multiplying the frequency of each phenotype after selection by the frequency of the cultural allele (which will not change between generations),

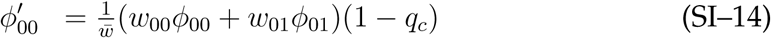

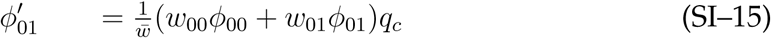

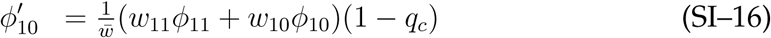

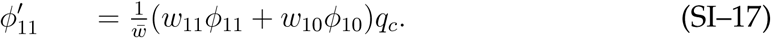

We wish to know when the mean population phenotype increases,

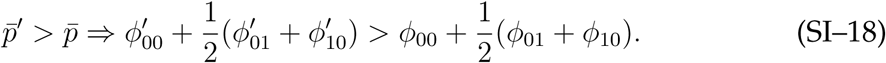

Substituting (SI–10a)–(SI–10d) and (SI–14)–(SI–17) into (SI–18), gives,

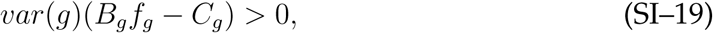

and multiplying both sides by 1 */var*(*g*) leads to (12).

### SI-4 Model 2

In this model, individuals encounter one another and play a prisoner’s dilemma. This time, the game determines both the reproductive fitness and cultural fitness of the players. We imagine individuals producing ‘cultural gametes’, or behavioral tokens. The probability of acquiring a given cultural allele will be determined by the proportion that allele constitutes of all the available cultural gametes. The expected number of cultural gametes produced by individuals of each type are:

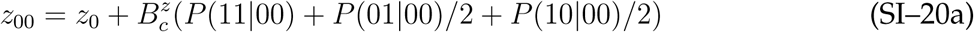

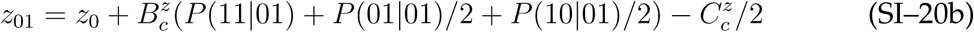

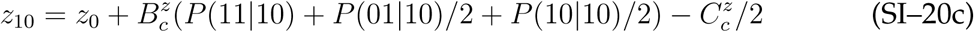

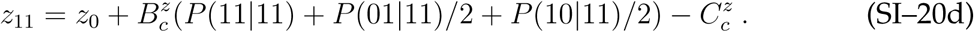

It is important to note that the terms 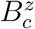 and 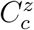 are the gametic fitness benefit and cost, as opposed to *B_c_* and *C_c_* that appear in (9). The actual cultural fitness of an individual *i* is 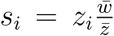. Writing the cultural fitnesses in terms of the *z_i_,* we can write the following recursions for the frequencies of types in this model:

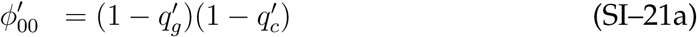

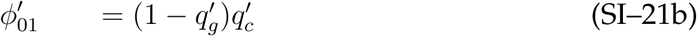

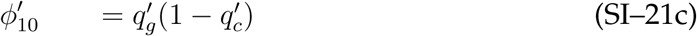

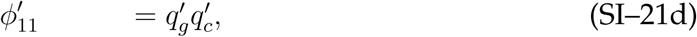

where,

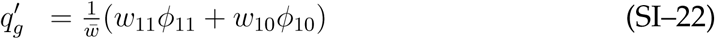

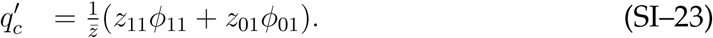

We wish to solve for the condition when the mean phenotype increases in the 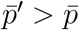,

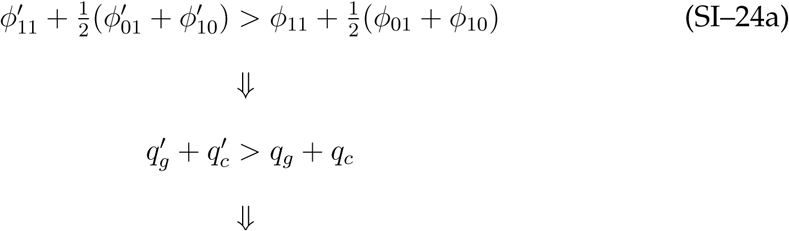

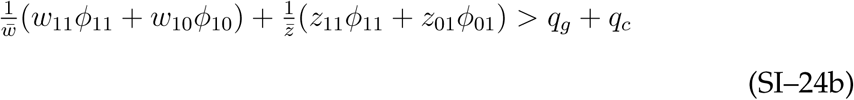

Substituting *q_g_* = *ϕ*_11_ + *ϕ*_10_, *q_c_* = *ϕ*_11_ + *ϕ*_01_, we can rewrite SI–24a as,

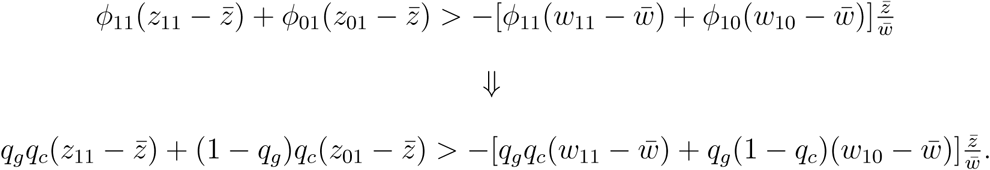

Finally, substituting the fitness expressions (SI-10a)-(SI-10d) into the RHS, and after considerable algebra, we arrive at,

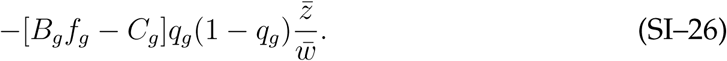

Similarly, substituting the (SI-20a) into the LHS, and after some further algebra, we have,

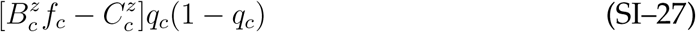

Combining the two sides of the inequality and dividing both sides by *q_c_*(1 – *q_c_*), we arrive at (14).

Alternatively, we could have arrived at (14) more directly by using (9). substituting for the cultural fitness, 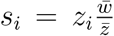, we can then rewrite (9) as,

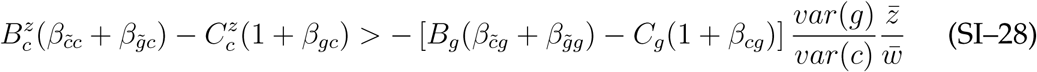

Again, we compute the relevant terms:

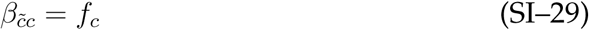

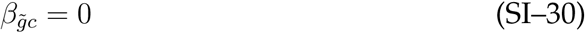

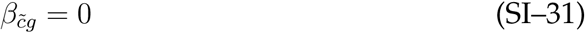

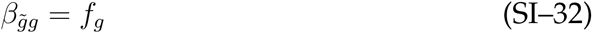

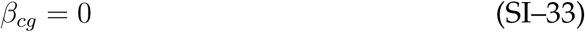

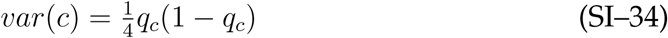

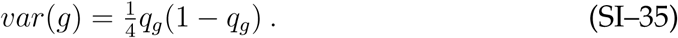

Substituting these terms into (9) gives us (14).

### SI-5 Covariance between genetic and cultural transmission

The previous models all assumed that offspring acquired their genotype and culture-type independently. Here we introduce a correlation between both types of inheritance. Let *k* be the probability that an offspring individual with certainty acquires the cultural allele that corresponds to their genetic allele (e.g. both genetic and cultural altruism allele). In this case we have the following encounter probabilities:

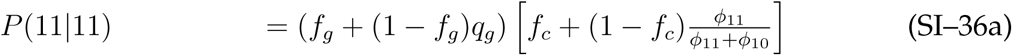

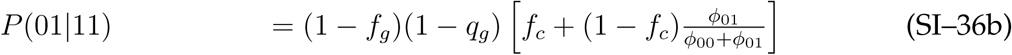

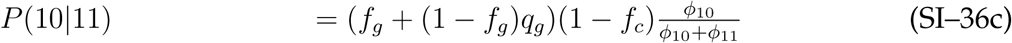

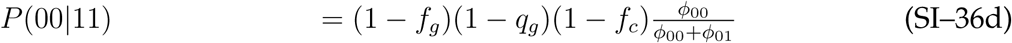

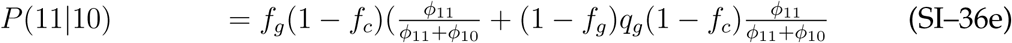

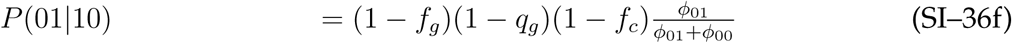

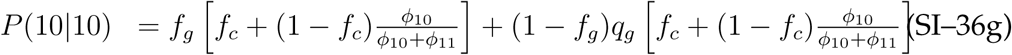

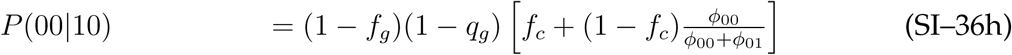

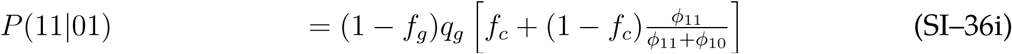

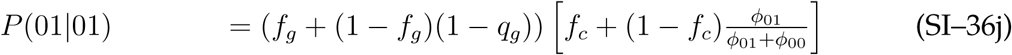

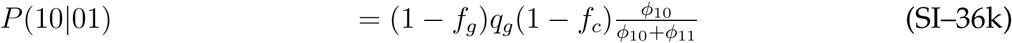

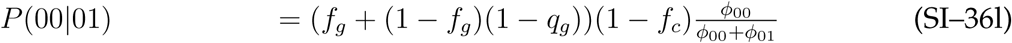

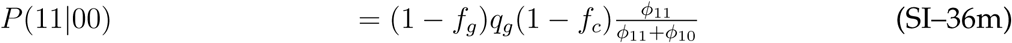

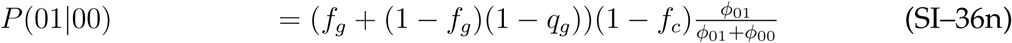

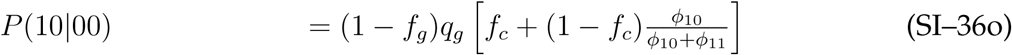

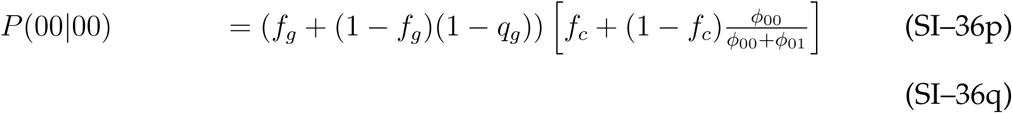

The recursions for the frequencies of types are:

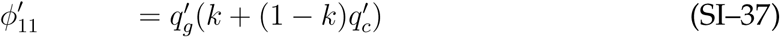

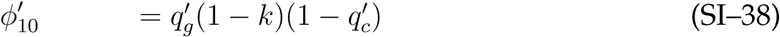

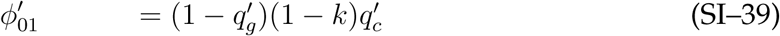

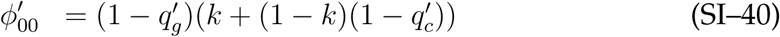

Our model occurs in two stages. First, each individual produces a number of genetic and cultural gametes according to selection in both domains. Then those gametes are paired, either according to gene-culture assortment, or the pure gamete frequencies. We will call the frequency of the altruistic alleles amongst gametes, *q_g_* and *q_c_*, and the frequency of the alleles amongst actual individuals, 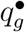 and 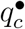.

Again, we consider the condition 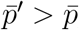, which leads to,

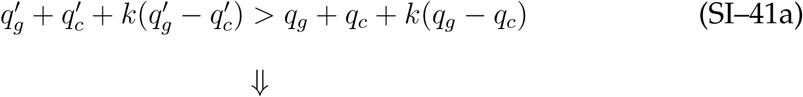

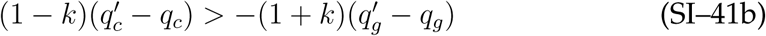

Here, we note that 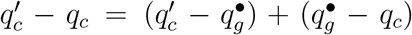. The first term is the change in frequency from ancestral individuals to descendant gametes, while the second term is the change in frequency from ancestral gametes to ancestral individuals.

We have,

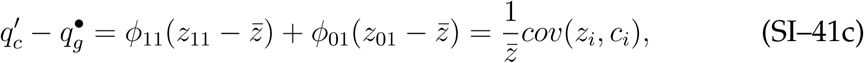

and,

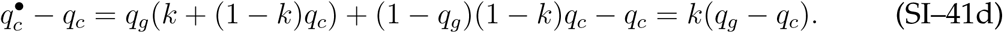

Noting that 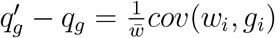, we have,

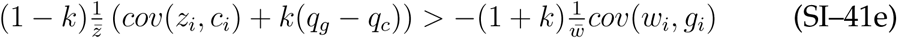

Expanding the covariance terms in the fitness functions and rearranging terms, gives,

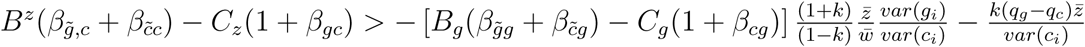

Computing all of the above regression coefficients we arrive at,

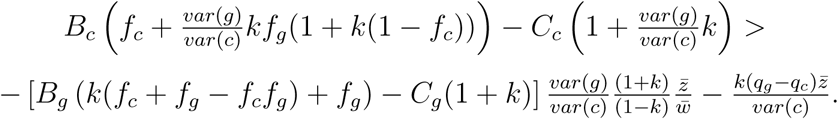

We can see that when *k =* 1—meaning genotype and culture-type are perfectly correlated—the condition is impossible to meet if genetic selection is opposed to altruism.

### SI-6 Non-additive phenotypes

We assume that all descendant individuals have a (potentially) unique function for mapping from heritable inputs to phenotype, *p_j_*(*f_j_*(*c*_1_, …, *c_N_*), *h_j_*(*g*_1_, ⋅, *g_N_*)). Assuming that the change in phenotype is small over small fluctuations in heritable inputs (e.g. because we are considering small evolutionary time scales), we can take a first order Taylor approximation of a phenotype function around the point (*c̄*, …, *c̄, ̄g*, …, *ḡ*) = (c̄, ḡ),

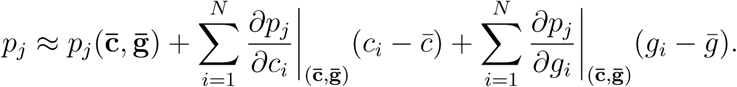

To obtain the Price equation, we can substitute the above expression into 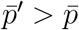,

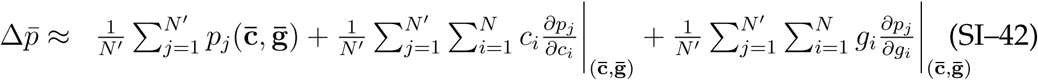

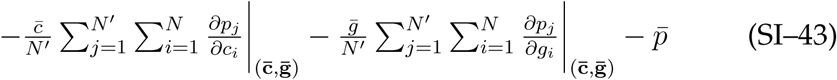

Switching the order of all the summations, and defining the quantities, 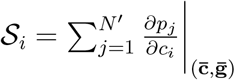 and 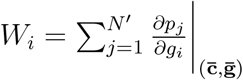, we can write,

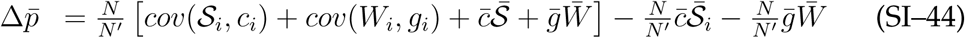

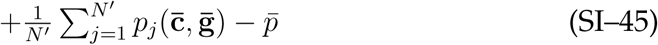

Canceling terms we arrive at Eq. (16).

If we continue our expansion of the phenotype function, we arrive at the following result,

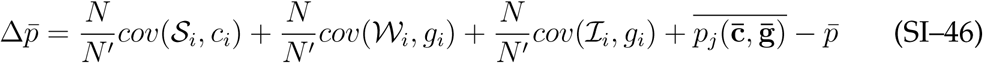

where,

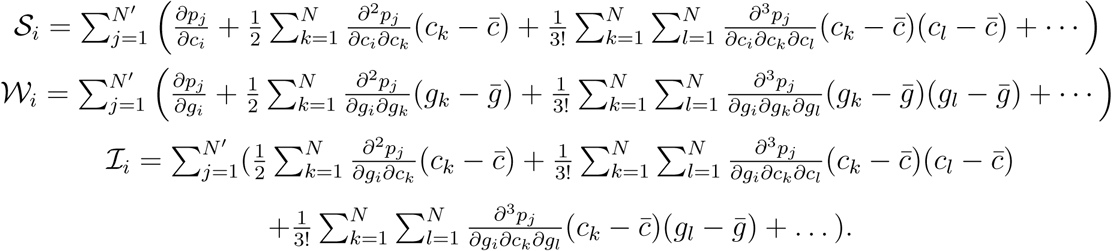

The dots represent higher order terms in the expansion. The *S_i_* and *W_i_* terms are exclusive to the cultural and genetic domains, while the *I_i_* term captures interactions between the two forms of inheritance. The additional covariance term captures the effect of interactions between genes and culture. In expanding these phenotype functions in a Taylor series, we’ve been able to directly relate the concepts of fitness to phenotype while making only minimal assumptions about either.

## Footnotes

1 In this derivation we assume that for every descendant *j* there exists some ancestor *i* for whom *γ_ij_* > 0.

